# Probing the chemical-biological relationship space with the Drug Target Explorer

**DOI:** 10.1101/308700

**Authors:** Robert Allaway, Salvatore La Rosa, Justin Guinney, Sara Gosline

## Abstract

Modern phenotypic high-throughput screens (HTS) present several challenges including identifying the target(s) that mediate the effect seen in the screen, characterizing ‘hits’ with a polypharmacologic target profile, and contextualizing screen data within the large potential space of drugs and biological screening model combinations. To address these challenges, we developed an interactive web application that enables exploration of the chemical-biological interaction space. Compound-target interaction data from public resources were quantified for over 280,000 molecules. Each molecule was annotated with a name and chemical structure, and every target was annotated with gene identifiers. The Drug-Target Explorer allows users to query molecules within this database of experimentally-derived and curated compound-target interactions and identify structurally similar molecules. It also enables network-based visualizations of the compound-target interaction space, and incorporates comparisons to publicly-available *in vitro* HTS datasets. Users can also identify compounds given one or more targets of interest. The Drug Target Explorer is a multifunctional platform for exploring chemical space as it relates to biological targets, and may be useful at several steps along the drug development pipeline including target discovery, structure-activity relationship, and lead compound identification studies.

## Introduction

In the modern drug discovery and development process, high-throughput screens (HTS) of drugs have become a common and important step in the identification of novel treatments for disease. In the past decade, studies describing or citing high throughput drug screening are increasingly prevalent, topping 1000 per year for the past 5 years (Figure 1) and span many disease domains such as cancer, neurodegenerative disease, and cardiopulmonary diseases. These screens are often phenotypic in nature whereby a large panel of compounds of known, presumed known, and/or unknown mechanisms of action are tested in a biological model of interest and generate phenotypic readouts such as apoptosis or proliferation. While these types of screens facilitate the rapid identification of biologically active drugs or chemical probes, they also present several challenges.

**Figure 1.**
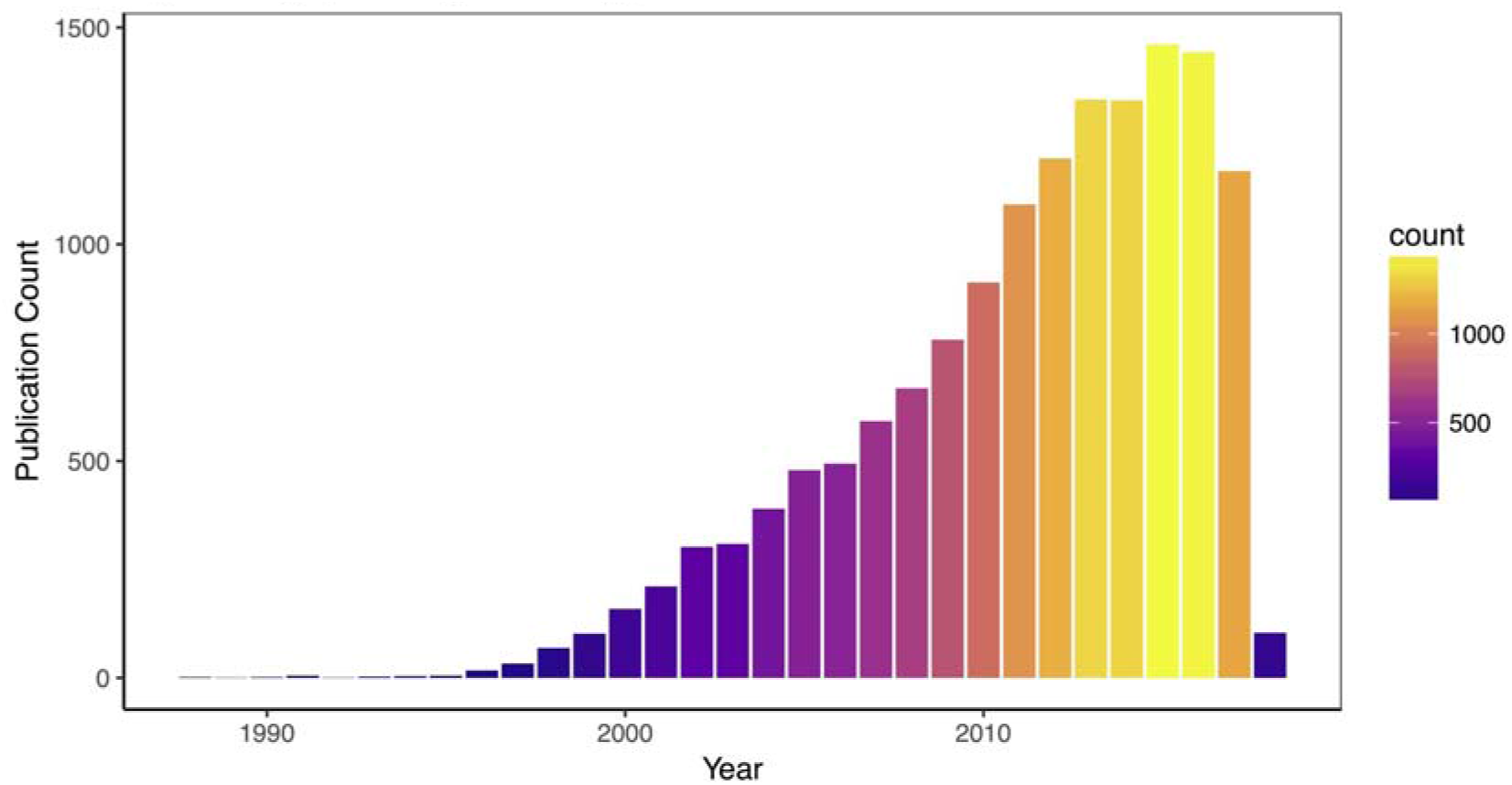
High throughput drug screening is an increasingly common experimental approach. Yearly count of Pubmed-indexed publications that appear with the search term “high throughput drug screening.” Search performed on January 30, 2018.

One prevailing challenge is the identification of the specific biological mechanisms within a cell that determine the response in a screen. The search for novel drugs constantly pushes the pharmaceutical researchers to include novel chemical sets in phenotypic screens, with the caveat that the underlying mechanism of action (MoA) of a particular compound cannot usually be gleaned from the phenotypic screens. (1) Most of the time, identifying the MoA requires additional experimentation, particularly if the molecule represents a novel or understudied chemical entity. Another challenge is that the polypharmacologic nature of many small molecules can make it difficult to interpret HTS results as a given drug may affect multiple targets with a range of efficacy. This, in turn, presents the difficulty of consolidating multiple targets into a unified biological mechanism or set of mechanisms leading to poorly annotated targets, misunderstood MoAs (2), and unknown or ambiguous off-targets with potential deadly side effects (3,4). A final challenge is that identification of related molecules and their targets is not always straightforward; in the context of HTS analysis, structurally and functionally related molecules that are not contained in a screening library might be useful to explore.

Multiple tools and databases have attempted to address various aspects of the challenges outlined above (see Table 1). These tools allow the user to explore known polypharmacology of small molecules. Many also allow users to explore compound-target relationships by querying either by molecule or by target: DGIdb, DT-Web, BindingDB, Polypharmacology Browser, STITCH, and SuperTarget allow users to identify MoAs/targets of a given compound by evaluating a query drug (5–10), while DT-Web, BindingDB, Polypharmacology Browser, and STITCH allow users to search by chemical similarity using any query molecule (Table 1). Probe Miner, alternatively, is designed primarily to handle target-based queries (11). All tools listed in Table 1 allow users to identify molecules with known polypharmacology, but only two, STITCH and SuperTarget, provide the ability to summarize these targets into biological pathways/mechanisms using a gene list enrichment approach (9,10). The final challenge - identifying structurally or functionally related molecules - is addressed by DT-Web, BindingDB, Polypharmacology Browser, and STITCH (6–9).

**Table 1.** Summary of selected features/uses of databases and applications for exploring molecule-target relationships and their overlapping features with the Drug-Target Database. Related tools include Probe Miner (11), DGIdb (5), DT-Web (6), BindingDB (7), Polypharmacology Browser (8), STITCH (9), ChEMBLSpace (12), and SuperTarget (10).

|  | Drug-Target Explorer | Probe Miner | DGIdb v3.0 | DT-Web | BindingDB | Polypharmacology Browser | STITCH | ChEMBLSpace | SuperTarget |
| --- | --- | --- | --- | --- | --- | --- | --- | --- | --- |
| Web app? | X | X | X | X | X | X | X |  | X |
| Open-source software? | X |  | X | X - underlying R package only | unknown | X |  | X |  |
| Search by targets to find drugs? | X | X | X | X | X | X - only by PDB-listed ligands | X | X | X |
| Search by drugs to find targets? | X |  | X | X | X | X | X |  | X |
| Identification of molecules that are associated with multiple query targets? | X |  |  | unknown |  |  |  |  |  |
| Drug structure input? | X |  |  | X | X | X | X |  |  |
| Drug name/ID input? | X |  | X | X | X | X | X |  | X |
| Visualize drug-target networks? | X |  |  | X, with user provided drug-target networks | not currently functioning |  | X |  |  |
| Identify chemically similar drugs? | X |  |  | X, with user provided drug-target networks | X | X | X |  | X |
| Allows queries using molecules not in database? | X |  |  | X, with user provided drug-target networks | X | X | X |  |  |
| Target organism? | human | human | human | human | human and others | human and others | human and others | unknown | human and others |
| Target space? | 3.6k | 2.2k | 6.1k | 3.8k | >7k | 4.6k | 9.6mil | unknown | >6k |
| Chemical space? | 280k | 400k | 10k | 4.4k | >642k | 870k | 500k | unknown | >196k |
| Quantitative interactions? | X | X |  | unknown | X | X | X | X | X |
| Qualitative interaction? | X | X | X | X |  |  | unknown |  | X |
| Explore polypharmacology? | X | X | X | X | X | X | X | X | X |
| Polypharmacologic target enrichment? | X |  |  |  |  |  | X |  | X |
| Comparison of query molecule to HTS drug response datasets? | X |  |  |  |  |  |  |  |  |
| Target prediction? |  |  |  | X | X | X |  |  |  |
| Database access? | Open | Open | Open | Open | Open | Open | Full database requires license | unknown | unknown |
| Last known update | 2018 | 2018 | 2018 | 2018 | 2018 | 2016 | 2016 | 2015 | 2012 |

While several of the tools listed address one or more of these challenges, there are some gaps (Table 1). For example, ChEMBLSpace does not have a web interface and therefore requires installation on a compatible system before use (12). In addition, not all of these tools are open-source (STITCH, SuperTarget, and BindingDB). An easy to modify open-source application could enable users to create features that are helpful for their specific analyses. While most tools allow both drug-based and target-based queries, none appear to facilitate queries for molecules that affect several targets, which may be useful for users who want to leverage polypharmacology by employing drugs that inhibit multiple biological mechanisms. While multiple targets can be queried at one time in STITCH, it is not straightforward to identify single molecules that affect all query targets. In addition, DGIdb and ChEMBLSpace cannot be used to explore similar chemical space to the query molecule. These two, plus SuperTarget, also cannot be queried using molecules that are not in the database; a feature that might help users with novel preclinical candidate drugs. With the exception of DT-Web and STITCH, these tools do not allow visualization of drug-target networks, which may help users address the challenge of identifying structurally or functionally related drugs. No tools other than STITCH perform gene list enrichment, which may help users interpret the biological MoAs of polypharmacologic molecules.

To address these gaps, we developed the Drug-Target Explorer. Specifically, the Drug-Target Explorer enables the user to (1) look up targets for individual molecules and groups of molecules, (2) explore networks of targets and drugs, (3) perform gene list enrichment of targets to assess target pathways of compounds, (4) compare query molecules to cancer cell line screening datasets, and (5) discover bioactive molecules using a query target and exploration of these networks. We anticipate that the users will include biologists and chemists involved in drug discovery who are interested in performing hypothesis generation of human targets for novel molecules, identifying off-targets for bioactive small molecules of interest, and exploring of the polypharmacologic nature of small molecules.

## Results

The Drug Target Explorer was designed to facilitate the following use-cases: hypothesis generation of targets for newly-discovered molecules, identification of off-targets for bioactive research molecules, and exploration of the polypharmacologic nature of many drugs. Below, we include vignettes highlighting how the Drug-Target Explorer can facilitate analysis in these areas.

### Identifying potential off-target effects of novel molecules

To highlight the use of this app to find potential off-targets of a novel molecule, we queried the Drug-Target Explorer for C21, a recently-published Polo like kinase (PLK) inhibitor that is not captured in our database (13). This molecule inhibits Plk2 and Plk1 in the low nM range, and Plk3 in the low uM range (13). Using a Tanimoto similarity of 0.65 or greater, we identified 14 molecules (Figure 2A, Supplemental Table 1). PLK1, PLK2, and PLK3 are known targets of several of these molecules, such as BI 2536 and volasertib. Curiously, CAMKK, BRD4, PDXK, and PTK2 are also targeted by molecules in this chemical set, with pChEMBL values >6-8. A plausible hypothesis could be that these targets are affected by this family of molecules, including the query molecule, in the 10-1000 nM range, which would indicate that further research is needed to determine the selectivity of C21 or other structurally related molecules.

**Figure 2.**
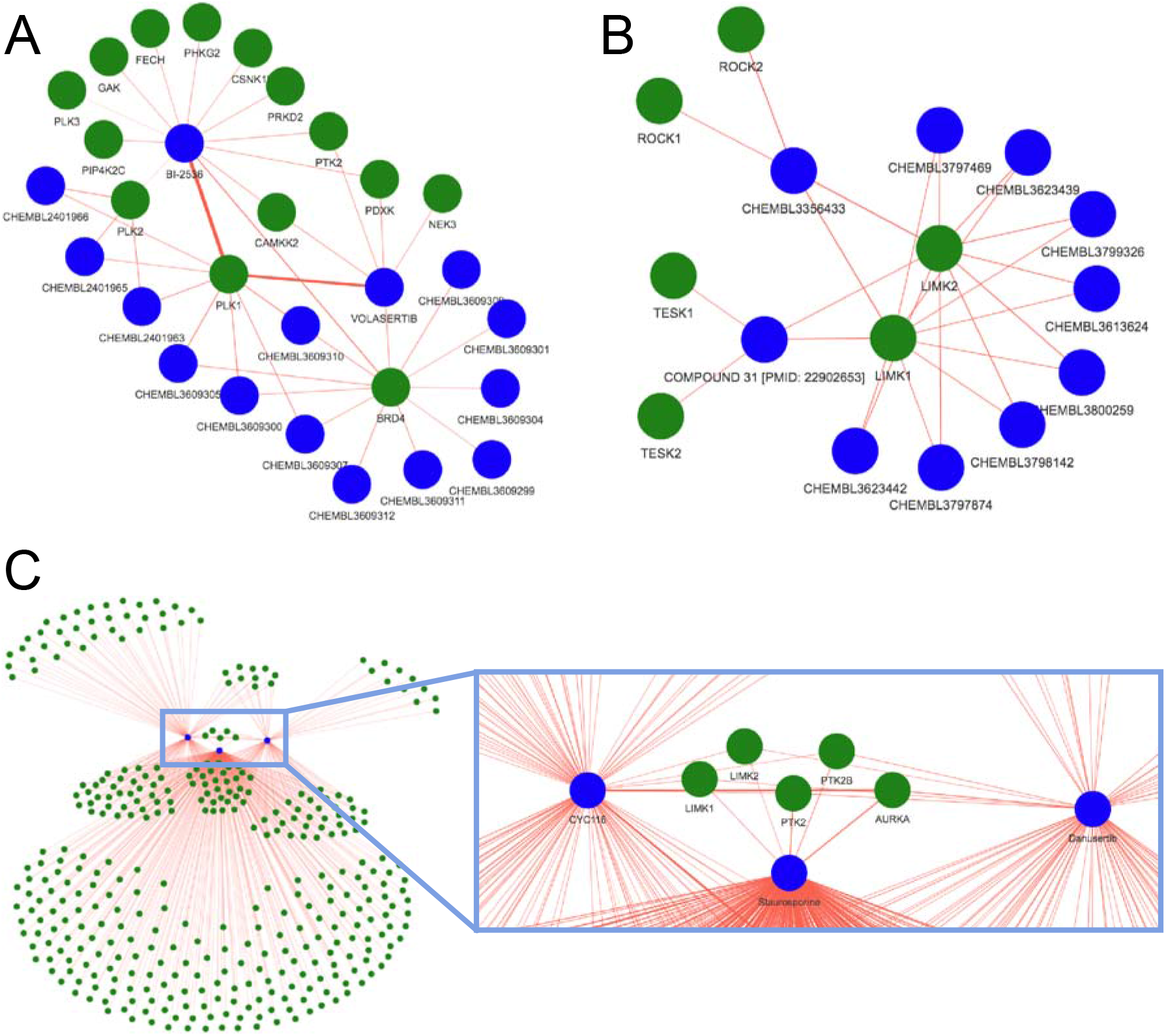
Molecule-target networks highlight targets within chemical families. (A) Using the novel Plk inhibitor C21 as a query with a Tanimoto cutoff of 0.65 (smiles: ccnc(=o)c1=cc2=c(c=c1)n(c=c2)c1=nc=c2n(c)c(=o)[c@@h](cc)n(c3cccc3)c2=n1), we identify 14 related molecules (blue vertices), and observe several targets (green vertices) common to multiple members of this family, including PLK1, PLK2, BRD4, CAMKK2, PTK2, and PDXK. (B) A gene-based query for two targets (green vertices), LIMK1 and LIMK2, identifies 10 molecules (blue vertices), as well as other targets affected by these molecules. (C) A query for multiple targets relevant to tumors caused by neurofibromatosis type 2 identifies three promiscuous molecules that have associations with these targets.

### Identifying off targets of existing molecules

This app may also be useful in identifying off-targets of existing molecules in a preclinical or exploratory research setting. In order to confidently interrogate the role of cellular targets, one must use compounds with specificity for those targets. A well-known example of a non-specific inhibitor is imatinib. This molecule, developed for use in the treatment of chronic myelogenous leukemia, was initially considered a selective inhibitor of Abl (14). More recently, several other targets have been identified for imatinib such as KIT, PDGFRA, and PDFGRB (15). Querying the Drug Target Explorer indicates that there is evidence for 61 targets of imatinib, several of which have pChEMBL values within a reasonable range of Abl, PDGFRA, and PDFGRB (Supplemental Table 2). These targets must all be considered when evaluating imatinib in human model systems.

A more recent example is the tool compound G-5555, a selective PAK1 inhibitor (16). This compound has been used to demonstrate the role of PAK1 in cellular processes such as invasion (17). A search of the Drug-Target Explorer database showed that this molecule not only binds PAK1 (mean pChEMBL = 8.01), but there is qualitative evidence for effects on PAK2/3, and quantitative evidence suggesting an effect on SIK2, MAP4K5, and PAK2 at similar concentrations of G-5555 (mean pChEMBLs 8.05, 8, and 7.96 respectively, Table 2). G-5555 also may have an effect on STK family proteins (STK3, STK24, STK25, STK26) and LCK. Therefore, any findings with G-5555 with regards to PAK1 inhibition must be validated with other selective inhibitors or genetic approaches, as Jeannott and colleagues did (using other PAK inhibitors such as FRAX597 and FRAX1036, as well as PAK1 silencing RNA), to confirm that the effects observed are PAK1 specific (17).

**Table 2.** Targets of G-5555 found in the Drug-Target Explorer Database.

| Molecule Name | HGNC Symbol | Mean pChEMBL | n Quantitative | n Qualitative | KSI | Confidence |
| --- | --- | --- | --- | --- | --- | --- |
| CHEMBL3770443 | PAK1 | 8.01 | 3 | 1 | 0.1 | 11 |
| CHEMBL3770443 | PAK2 | 7.96 | 2 | 1 | 0.1 | 9.96 |
| CHEMBL3770443 | SIK2 | 8.05 | 1 |  | 0.1 | 9.05 |
| CHEMBL3770443 | MAP4K5 | 8 | 1 |  | 0.1 | 9 |
| CHEMBL3770443 | STK26 | 7.7 | 1 |  | 0.1 | 8.7 |
| CHEMBL3770443 | STK25 | 7.47 | 1 |  | 0.1 | 8.47 |
| CHEMBL3770443 | STK24 | 7.37 | 1 |  | 0.1 | 8.37 |
| CHEMBL3770443 | STK3 | 7.37 | 1 |  | 0.1 | 8.37 |
| CHEMBL3770443 | LCK | 7.28 | 1 |  | 0.1 | 8.28 |
| CHEMBL3770443 | PAK3 |  |  | 1 | 0.1 | 1 |

### Identifying polypharmacologically-targeted pathways and drugs with similar biological effects

In order to provide biological context, this app allows the user to aggregate multiple targets from compounds into functional categories. Using the previous example of G-5555, we performed enrichment analysis on the list of targets to identify potential biological pathways and MoAs that this molecule may disrupt. In doing so, we observed that G-5555 targets are enriched in several Gene Ontology terms and KEGG Pathways like T-cell receptor signaling, Ras/MAPK signaling, and Golgi-localized proteins (Supplemental Table 3). The app also allows the user to compare the query molecule to drugs in the Cancer Cell Line/CTRP and GDSC/Sanger cell line screening datasets. Specifically, the app identifies the most similar molecule available in these datasets and uses that molecule as a reference to plot chemical similarity vs drug response correlation.

### Finding a drug for known targets

Finally, the tool allows users to perform a reverse search, i.e. identify molecules that have an association with a query target or targets and assess the known selectivity of these molecules. For example, Petrilli et. al. identified LIM domain kinases as targets of interest in tumors caused by the genetic disease neurofibromatosis type 2 (NF2) (18). They found that pharmacologic (LIMK1/2 inhibitor BMS-5) and genetic modulation of LIMK1 and LIMK2 caused cell-cycle inhibition and reduced viability in merlin *(Nf2)* deficient Schwann cells (18). In the context of follow-up and validation studies, it may be beneficial to use alternate molecules that target LIMK1/2 at the same or greater potency than BMS-5. We used the Drug-Target Explorer to find molecules that target LIMK1 and LIMK2 (Supplemental Table 4, Figure 2B). For example, BMS-5 (CHEMBL2141887 in the Drug-Target Explorer) has mean pChEMBLs of 7.33 and 7.07 for LIMK1 and LIMK2 respectively. A good alternative to validate the effects of this molecule might be CHEMBL3623442, a relatively structurally distinct small molecule (extended fingerprint Tanimoto similarity of 0.433 to BMS-5 in this database), with pChEMBLs of 9 and 8.52 for LIMK1 and LIMK2 respectively. Another interesting possibility is the identification of multiple molecules with overlapping desired targets and non-overlapping off-targets to reduce off-target effects, or to identify synergistic/additive single-target, multi-drug combinations as outlined by Fitzgerald et al 2006 (19). Using the above scenario with LIMK1/2, it may be possible to use structurally distinct molecules in combination or in sequence, like CHEMBL3356433 and Compound 31 highlighted in Figure 2B, to reduce off-target effects or inhibit LIMK1/2 in an additive or synergistic manner. The opposite approach could also be taken by finding a single molecule that binds multiple desired targets. In the case of merlin-deficient cells, focal adhesion kinases (FAKs) such as PTK2 (FAK2) and PTK2B, as well as Aurora kinase A (AURKA) have been highlighted as potential targets of interest (18,20,21). Using the Drug-Target Explorer, we can identify molecules that target LIMK1/2, PTK2/2B, and AURKA (Supplemental Table 5, Figure 2C). Using this information, a rational hypothesis might be that CYC116 or danusertib could be effective and selective for NF2-deficient tumor cells; to our knowledge, the use of these molecules in this setting has yet not been explored.

## Discussion

In the present study, we demonstrate that the Drug-Target Explorer enables the user to look up targets for novel and known molecules such as C21, G-5555, and imatinib, as well as explore networks of these drugs and their targets. Users can perform target enrichment to consolidate multiple targets to into pathways, compare query molecules to screening datasets, and identify bioactive molecules given a query target.

Several future directions are envisioned for this application. The code and database has been designed in such a way that any database with structural information and drug-gene target information (qualitative associations, or quantitative associations that can be coerced to pChEMBL values) can be harmonized and integrated into the database. Therefore, as new datasets become available, such as the recently-published Drug-Target Commons (22), they can be integrated and released. We also envision occasional errors being identified as the database is explored and vetted by users and have included a feedback form for users to suggest new data to integrate, as well as to highlight necessary corrections to the dataset. Currently, the query molecule to full database similarity calculation is computationally intensive. One solution to speed up calculation times may be to implement a locality sensitive hashing method in future versions of the database and web app, such as the method devised by Cao et al 2010 (23). An additional planned feature for this app is the implementation of a bulk annotation feature to allow users to annotate HTS data with targets and/or putative targets of identical or structurally related molecules. Finally, the integration of a predictive framework for identifying targets of query drugs based on drug and target feature data would enable users to quantitatively predict targets of novel molecular entities rather than manually exploring structurally similar molecules.

The Drug-Target Explorer enables users to explore known molecule-human target relationships as they relate to chemical similarity rapidly and with minimal effort. We anticipate that users such as biologists and chemists using chemical probes or studying preclinical therapeutics will find this tool useful in several areas. Specifically, this tool may aid drug discovery efforts by accelerating hypothesis generation, simplifying the transition from phenotypic HTS results to mechanistic studies, and streamlining the identification of candidate molecules that target a protein or mechanism of interest.

## Methods

To build the database of known compound-target interactions, we aggregated five data sources containing qualitative and quantitative interactions (Figure 3). We considered qualitative interactions to be curated compound-target associations with no associated numeric value. Quantitative interactions were defined as compound-target information with a numeric value indicating potency of compound-target binding or functional changes. Qualitative compound-target associations were retrieved from the DrugBank 5.0.11 XML database, the DGIdb v3.0.1 interactions.tsv file, and ChemicalProbes.org (acc. Jan 17 2018) (5,24,25). pChEMBL, IC50, C50, EC50, AC50, Ki, Kd, and potency values for *Homo sapiens* targets were retrieved from the ChEMBL v23 MySQL database (26). Kd values were also obtained from Klaeger et al 2017, in which the authors determined the Kd of 244 kinase inhibitors against 343 kinases (27). For all quantitative and qualitative data sources, compound structural information (SMILES) was retrieved when available. When not available, it was batch annotated using the Pubchem Identifier Exchange Service, or, in some cases, manually annotated via PubChem and ChemSpider search (28,29).

**Figure 3.**
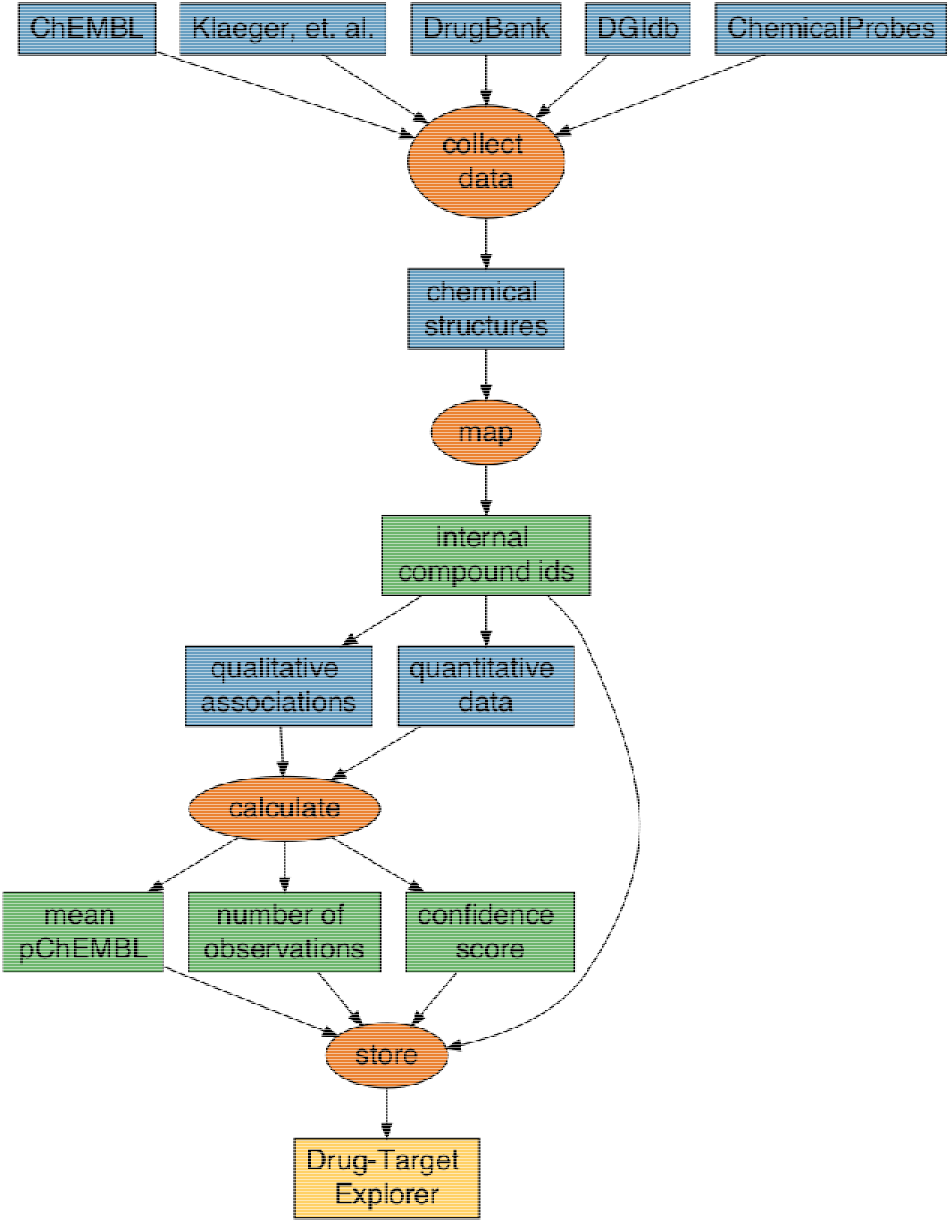
Process for developing the Drug-Target Explorer. Molecule-target and chemical structure data were collected from public sources. In the case of DGIdb, chemical structures were assigned using the PubChem Chemical Identifier Exchange or manually assigned using ChemSpider and PubChem. Chemical structures were converted to circular fingerprints and the databases were mapped to internal Drug-Target Explorer identifiers. Qualitative and quantitative data were summarized by calculating several summary statistics, and these data were stored together with the internal identifiers to form the Drug-Target Explorer database.

To consolidate data for “identical” molecules within and across multiple databases, the functional connectivity fingerprint (FCFP6)-like ‘circular’ fingerprint for each SMILES was calculated using the R interface (rcdk) to the Java Chemical Development Kit (CDK) (30–32). The package was modified to use the latest version of the CDK (2.1.1), which enables perception of chiral centers, enabling differentiation between isomeric molecules. Each unique circular fingerprint and all external IDs and SMILES associated with that fingerprint were then assigned an internal identifier, so that groups of molecules with identical fingerprints were assigned to the same internal ID. The internal molecular IDs were then mapped to each database to permit their aggregation. All datasets were combined and summaries were generated for each compound-target comparison using functions from the R ‘tidyverse’ (33).

The summary metrics described in Table 3 were calculated. One of these metrics, pChEMBL, is used to convey the efficacy of a given molecule. It is calculated from one of several semi-comparable values in the ChEMBL database, and is defined as the negative log 10 molar of the IC50, XC50, EC50, AC50, Ki, Kd, or potency (26). For example, a pChEMBL value of 7 would indicate that there is a measurable effect on a given target in the presence of 100 nM of molecule. To harmonize the data from Klaeger et al with ChEMBL data, the Kd values were converted to pChEMBLs. The mean pChEMBL was calculated for every molecule-target combination, as well as the number of quantitative and qualitative associations found in the source databases.

**Table 3.**
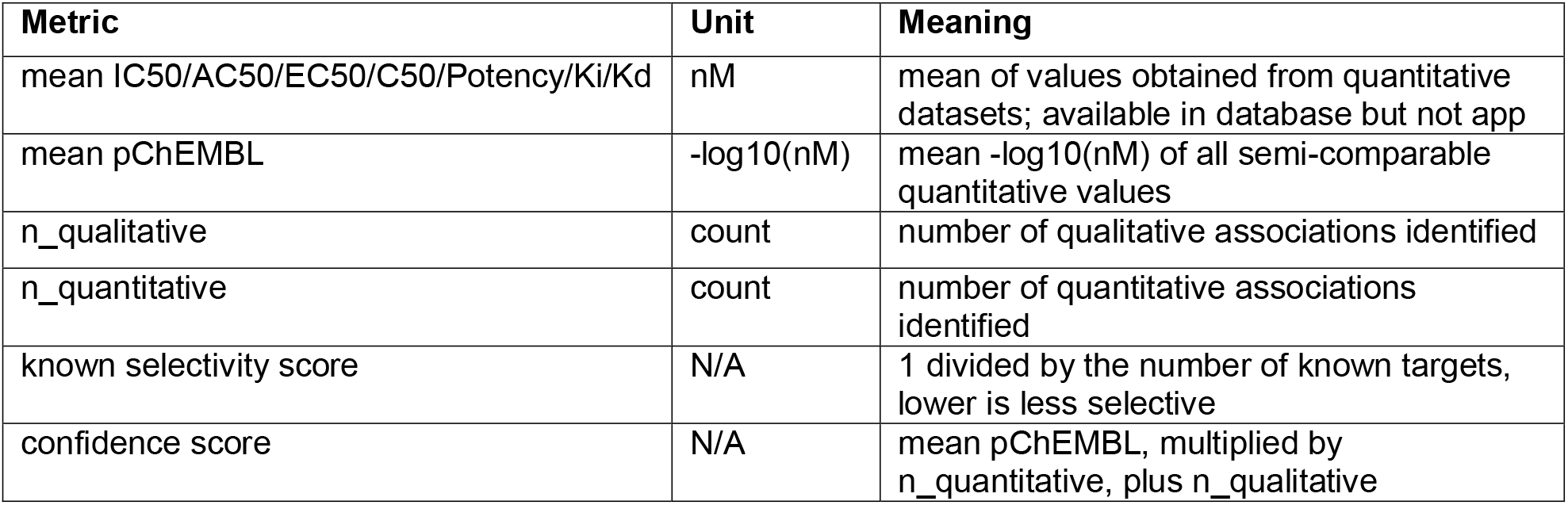
Drug-target association metrics summarized in the Drug-Target Explorer database.

| Metric | Unit | Meaning |
| --- | --- | --- |
| mean IC50/AC50/EC50/C50/Potency/Ki/Kd | nM | mean of values obtained from quantitative datasets; available in database but not app |
| mean pChEMBL | -log10(nM) | mean -log10(nM) of all semi-comparable quantitative values |
| n_qualitative | count | number of qualitative associations identified |
| n_quantitative | count | number of quantitative associations identified |
| known selectivity score | N/A | 1 divided by the number of known targets, lower is less selective |
| confidence score | N/A | mean pChEMBL, multiplied by n_quantitative, plus n_qualitative |

We calculated a known selectivity score for each molecule, which we defined as 1 divided by the total number of targets for that molecule (lower values correspond to lower molecule selectivity), and a confidence score for each molecule-target relationship, which we defined as the mean pChEMBL multiplied by the number of quantitative measurements, in addition to the number of qualitative annotations. A larger confidence score indicates greater confidence in this relationship; this confidence is weighted by the potency to give increased preference to high-potency compound-target interactions.

This resulted in a database containing 3645 human targets (represented by HUGO gene symbols), ~280,000 small molecules, and ~623,500 molecule-target relationships summarized from ~598,000 quantitative associations and ~25,000 qualitative associations. Finally, this database as well as fingerprints and chemical aliases for each molecule were saved as R binary files and stored on Synapse. All of the data, as well as snapshots of the source databases used to build the Drug Target Explorer database (with the exception of DrugBank, which requires a license to access) are accessible at www.synapse.org/dtexplorer. The Drug-Target Database is licensed under CC BY-SA 4.0.

We developed a Shiny application to permit exploration of the database (34,35). For chemical queries, users can search for molecules in the database by one of three methods: from a list of aliases obtained from the source databases, retrieving the chemical structure using the ‘webchem’ interface to the Chemical Identifier Resolver, or by directly inputting the SMILES string (36). A Tanimoto similarity threshold allows the user to narrow or widen the chemical space of the results. After querying, the input molecule is converted to a fingerprint and it’s similarity calculated relative to all molecules in the database, using ‘extended’ fingerprints. The user then can view the resulting set of molecules as well as the molecule-target relationships in interactive tables and graphs (Figure 4). In addition, the user can remove or include molecules on an a-la-carte basis, view the 2D structural representation of the input molecule, and perform target list enrichment analysis (37,38). Furthermore, the query molecule can be compared against molecules in the CTRP and Sanger cancer cell line drug-screening datasets to identify identical or similar structures in these datasets, and compare the relationship between chemical structure and correlations in drug response.

**Figure 4.**
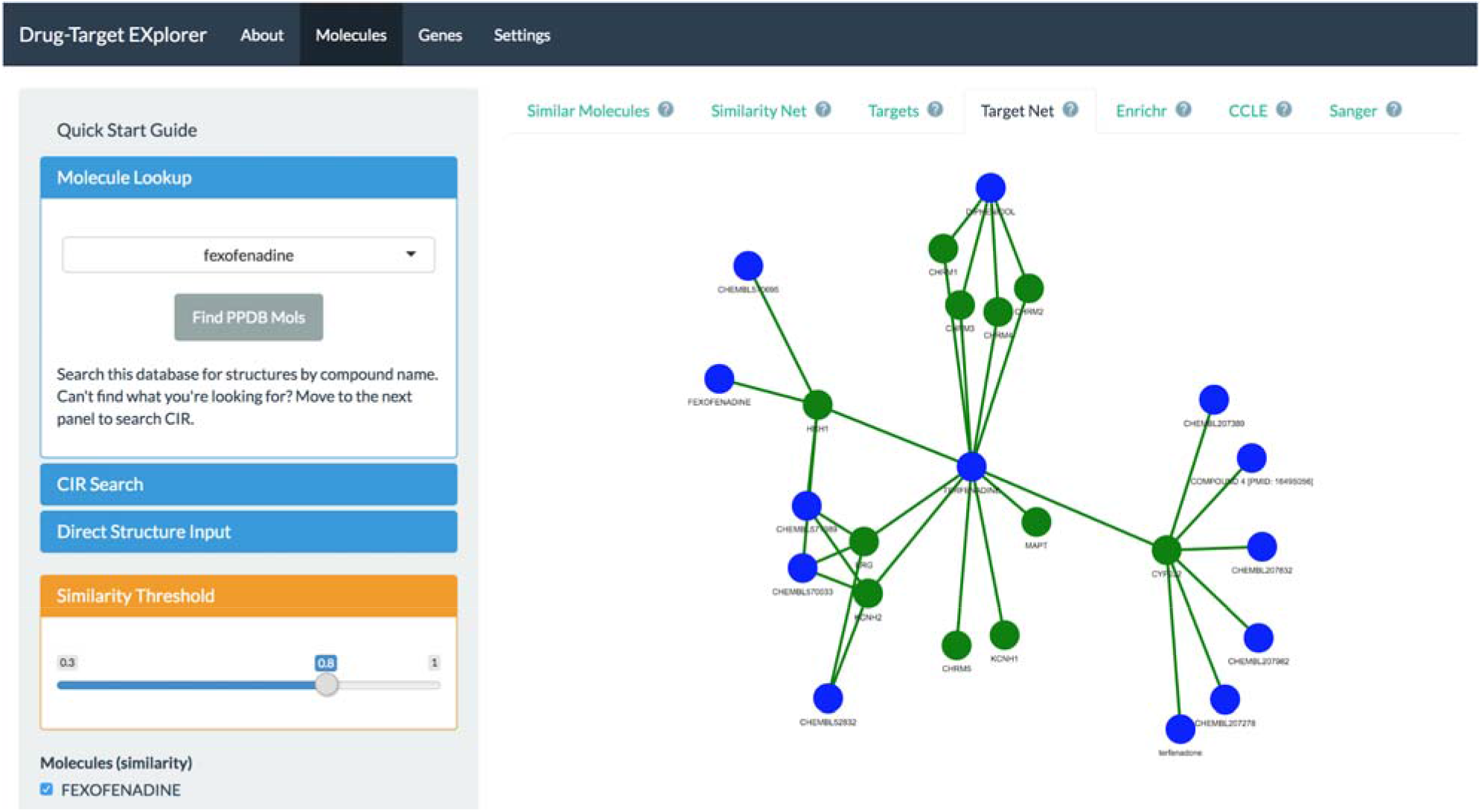
Layout of the Drug-Target Explorer. The “About” tab describes the apps functions and uses, the “Molecules” tab permits molecule-based searching, the “Genes” tab permits target queries, and the “Settings” tab allows the user to pick the fingerprinting method used.

For target queries, users can input one or more query HUGO gene(s) and identify molecules that are reported to bind those targets, and view these data in an interactive table. Users can also view these drugs in an interactive graph format to view their association with the query target and their other targets. The Drug-Target Explorer is available at www.synapse.org/dtexplorer. The source code for the Drug-Target Explorer app is available at https://github.com/Sage-Bionetworks/polypharmacology-db. The source code is licensed under Apache 2.0.

## Supplemental table legends

**Supplemental Table 1 – Targets of C21-like compounds in the Drug-Target Explorer Database**.

**Supplemental Table 2 – Targets of imatinib in the Drug-Target Explorer Database**.

**Supplemental Table 3 – Target enrichment analysis of G-5555 highlights putative mechanistic effects**. G-5555 targets were enriched in multiple Gene Ontology terms and KEGG pathways.

**Supplemental Table 4 – Molecules targeting LIMK1/2**. The database was queried for molecules that may modulate LIMK1 and LIMK2; this analysis revealed a large set of putative tool compounds.

**Supplemental Table 5 – Identification of multi-kinase-targeting molecules for NF2**. A query of the database for molecules that target several kinases of interest in NF2 (AURKA, LIMK1/2, PTK2/2B) identified 3 polypharmacologic compounds.

